# Mesoscale, Cantilever-Based Indentation Device for Mechanical Characterization of Soft Matter and Biological Tissue

**DOI:** 10.1101/758342

**Authors:** Andres Rubiano, Chelsey S. Simmons

**Affiliations:** Department of Mechanical and Aerospace Engineering, Herbert Wertheim College of Engineering, University of Florida, Gainesville, FL, USA; J. Crayton Pruitt Family Department of Biomedical Engineering, Herbert Wertheim College of Engineering, University of Florida, Gainesville, FL, USA; Division of Cardiovascular Medicine, College of Medicine, University of Florida, Gainesville, FL, USA

## Abstract

Tissue engineering has been driving a growing interest in mesoscale tissue mechanics (10^−4^ – 10^−2^ m), requiring tools to compare modulus between irregularly shaped primary tissue explants and synthetic scaffolds. We have designed and built a simple cantilever-based mesoscale indentation device to record force-displacement data during spring-loading, stress-relaxation, and creep experiments. Its simple design enables quantification of a wide range of soft matter moduli, from ~500 Pa collagen hydrogels to ~2 MPa silicones, by its compatibility with cantilevers of different stiffnesses and indentation probes of different sizes. A piezo-electric stage is used to drive a cylindrical or spherical indentation tip into the sample, while custom programming in LabVIEW through a data acquisition card enables stage control and acquisition of cantilever deflection using a capacitive sensor. Cantilever stiffness, deflection, and piezoelectric stage positions, acquired at a rate of 10Hz, are used to calculate force and indentation depth throughout indentation cycles. Using *xyz* manual coarse stages, tissue properties can be mapped across the sample surface. We have also built in commands to tune initial tip location using the piezo-stage to more easily find the sample surface, which is critical for accurate application of contact models. Here, we provide detailed information on how to design, build, and code a system for mesoscale indentation.

## I. INTRODUCTION

Understanding and controlling the mechanical properties of tissue is an increasingly important part of regenerative medicine and tissue engineering research. Cells can convert mechanical signals into biochemical signals, which can alter their morphology, proliferation, migration, differentiation, gene expression, protein production, and cell death. Mechanical properties of engineered constructs thus affect cell behavior, wound healing, and implant integration, while utilization of different cell populations and scaffold materials can affect mechanical performance of engineered cell-populated scaffolds and remodeled tissues. The reader is directed to *Mechanobiology of Cell-Cell and Cell-Matrix Interactions* edited by Wagoner Johnson and Harley for in depth reviews of these topics.^1^

Numerous mechanical characterization techniques have been adapted to tissue engineering applications, such as tensile testing, nanoindentation, and atomic force microscopy (AFM). However, for biological tissue and compliant biomaterials, apparent elastic moduli can vary across three orders of magnitude for a single tissue type depending on characterization technique and testing parameters.^2,3^ AFM is often used for mechanical characterization of biological tissue and other soft matter at the ~100 nm – 10 μm scale.^4^ Though extensive work has been put into creating standardized procedures,^5^ nano/microscale indentation is problematic for tissue-scale characterization as nano/microscale indentations quantify properties of a very specific region of a cell, e.g. nucleus, or one of the many components of the extracellular matrix. Bulk tissue properties do not match their nano/microscale counterparts, and even minor changes in indenter dimensions can create noticeable differences in obtained values.^6^ Additionally, for the commonly used Hertz contact model, ^7^ nano/microscale indentation aggravates the violation of its sample-homogeneity assumption, as confirmed by finite element simulations studies.^8^ Finally, indentation curves are highly strain rate dependent (Supplementary Figure A), which limits comparison between tissue and synthetic scaffolds across research groups: effective moduli will vary if there was any difference in indentation rate, tip diameter, contact model, and/or constitutive model used. Rheology is often used instead to characterize time-dependent mechanical properties of tissue, but rheological equipment and standard sample geometries are not easily applicable to arbitrarily shaped tissue resections.

Mesoscale indentation (10^−4^ – 10^−2^ m) is advantageous for measuring, comparing, and matching tissue mechanics because: a) indentation contact radii are on the order of “bulk” properties of tissue rather than cell-level heterogeneity, b) indentation parameters, e.g. sample size, indentation rate, indentation depth, and relaxation times, can be matched between biological tissue resections and synthetic scaffolds for accurate comparison, and c) obtained force-displacement data can be paired with complex constitutive models appropriate for computational methods or simple constitutive models to obtain consistent, comparable metrics between samples, experiments, and labs.

Our custom MesoScale Indentation (MSI) system supports mechanical characterization via mesoscale indentation, and MSI can accommodate a range of sample stiffnesses by easily switching components. Indentation probes can be effortlessly swapped to compensate for sample heterogeneity and modulus, and cantilevers with different design parameters can be used to adapt to the force range we expect from a given sample. This system is capable of outputting simple force-displacement data that can be easily processed with contact and constitutive models that best suit the tissue or sample type. Because components are controlled by custom LabVIEW code, raw force-displacement data can be analyzed in ways appropriate to soft matter rather than relying on commercial code often designed for rigid materials.

Additionally, our equipment offers simplicity in sample positioning and surface finding. We have added the ability to control the initial position before starting an indentation cycle to more accurately find sample surface, which is critical to fitting contact models.^9,10^ Finally, the system’s compatibility with a USB microscope (Dino-Lite Edge AM4115T-GFBW) underneath the transparent indentation surface enables monitoring of deformation in real-time. This manuscript details the design, construction, and coding of this custom, cantilever-based, mesoscale indentation system that can be used to quantify and compare mechanics of soft matter (~500 Pa – 2 MPa), both biological and synthetic. We also include procedures for part calibration, calculations of error propagation, and analysis of example data.

## II. APPARATUS AND PRINCIPLE OF OPERATION

A piezo-electric stage (P-628.1CD, Physik Instrumente) is controlled through a piezo-amplifier (E-625.CR Servo Controller, Physik Instrumente) to vertically move an aluminum plate to which a cantilever and capacitive sensor arrangement have been attached (Figure 1C-2). The capacitive sensor (C8S-3.2-2.0 and C8S-2.0-2.0, Lion Precision, Figure 1C-5) is mounted to a manual 25 x 25 mm aluminum translational stage (Opto-sigma, TAM-251S STAGE, Figure 1C-3) by a rectangular aluminum beam (Figure 1C-4). The small translational stage is used to bring the sensor near the top surface of the upper titanium flexure of the cantilever, within the sensor’s range of operation.

**Figure 1.**
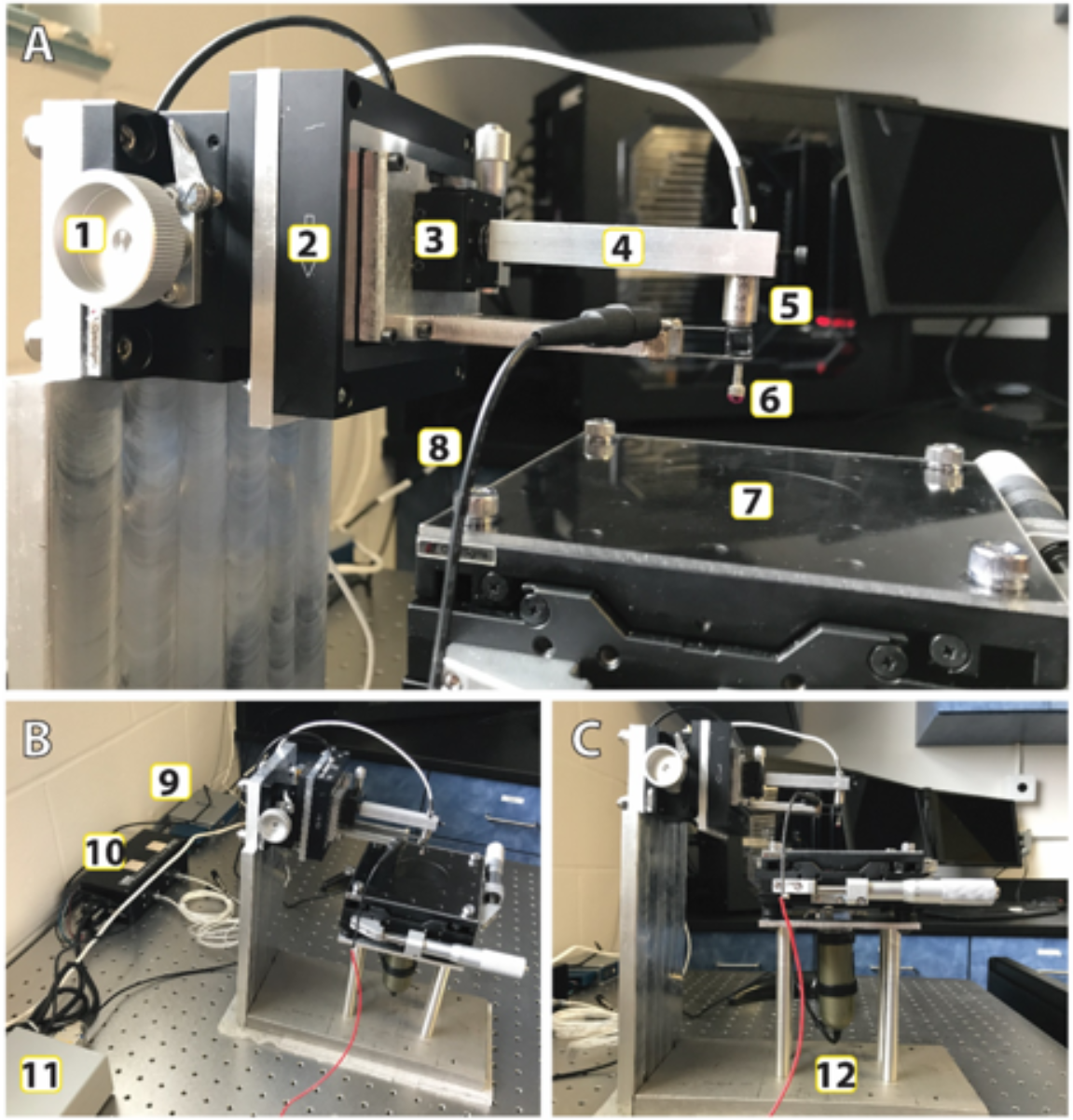
(A) Close-up of moving components: [1] manual coarse z-stage, [2] vertically positioned piezo-electric x-stage, [3] manual z-stage, [4] rectangular aluminum beam holder for [5] capacitive sensor, [6] indentation probe, [7] manual xy-stage with transparent acrylic surface, and [8] grounded wire connected to cantilever. (B) View of indentation device with all of its components: [9] Piezo-electric stage servo controller, [10] capacitive sensor compact driver, and [11] USB DAQ module for communication with controlling PC. (C) Sideview of indentation device showing [12] USB microscope.

The piezo-electric stage drives both the indentation probe (Figure 1C-10) – located at the free end of the cantilever – and the capacitive sensor, at the same rate. When the indentation probe comes into contact with the indentation sample and the stage is moved downward, the sample causes the cantilever to bend upward relative to the rest of the piezo-electric stage structure, which reduces the gap between the edge of the cantilever’s top surface and the capacitive sensor.

Cantilever deflection is calculated from the sensor-detected change in voltage, and the voltage-to-distance relationship of the capacitive sensor (e.g. our C8S-3.2-2.0 has a range of 500 μm, measured over 20.41 V). The capacitive sensor can be easily replaced to adjust resolution or range. Cantilever stiffness and deflection are used to calculate the interaction force between the probe and the sample, usually referred to as indentation force.

Indentation depth, i.e. the relative displacement of the cantilever tip or depth the probe travels inside the indented sample, is calculated as:

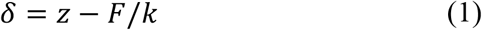

Where *δ* is indentation depth, *z* is piezo-stage displacement, *F* is indentation force, and *k* is cantilever stiffness.

LabVIEW is used to control the indentation profile and acquire time, stage displacement, indentation depth, and indentation force data. Our spring-loading experiments contain three automated phases: loading, holding, and retraction, though any open- or closed-loop waveform could be programmed. In the loading phase, the probe starts at the exact point where it first makes contact with the sample. The stage is moved a specified distance either in a defined time (loading time, *t_l_*) or using a specified indentation rate, 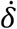 (due to very small cantilever deflections, stage displacement rate 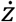, is assumed to be close to 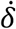). During the holding or relaxation phase, the probe is maintained in the same position for a specified time (relaxation time, *t_R_*), during which the sample is allowed to undergo stress-relaxation. During the spring-loading mode, the position of the piezo-stage is held constant, not the indentation depth; however, closed-loop control is possible in the LabVIEW program. For most of our samples, the indentation depth changed by less than 5% of the overall depth during relaxation, so we chose to utilize the spring-loading experiment rather than re-introduce additional noise from the piezostage to enforce closed-loop control of displacement. The final phase is the unloading phase, where the piezo-stage goes back to its initial position in the user-specified time (*t_u_*).

The program outputs a *.csv spreadsheet with time, piezo-stage displacement, indentation depth, and indentation force in column arrays. This information can be used to calculate desired mechanical properties using various contact and constitutive models of interest and relevance.

### II.A. LabVIEW Input/Output and Controls

LabVIEW was used to write a custom algorithm (Figure 2) that allows piezo-electric stage control and acquisition of cantilever deflection using a capacitive sensor (C8S-3.2-2.0 and compact driver CD1-CD6, Lion Precision) through a data acquisition card (NI 9220 and cDAQ-9171), National Instruments).

**Figure 2.**
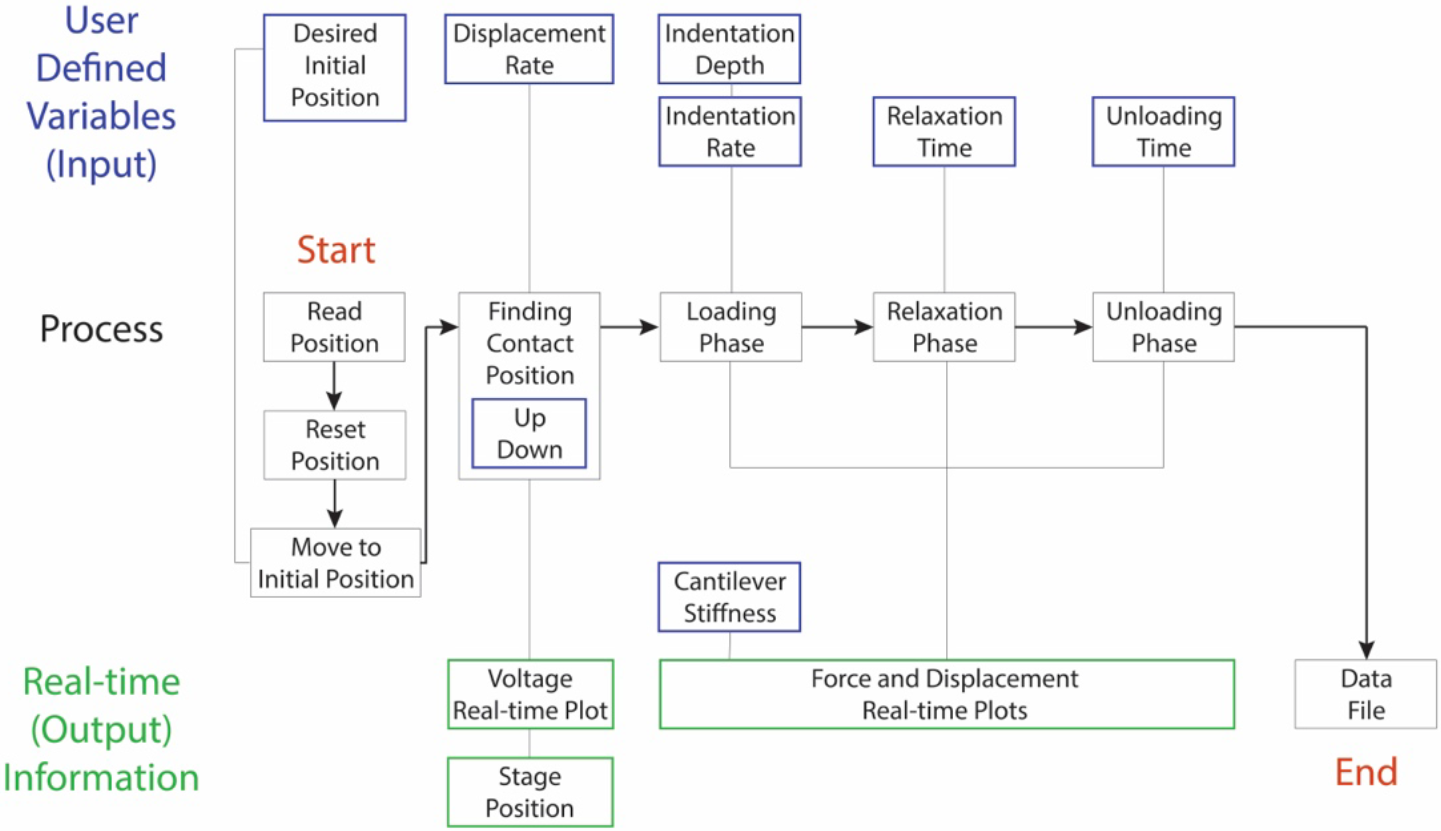
Process diagram of indentation cycle. The user-interface input controls for the indentation cycle include indentation depth, indentation rate, relaxation time, and unloading time. Additional controls include cantilever stiffness and indentation probe radius inputs – used for contact model calculations – as well as sample name and indentation number inputs, used for naming raw-data files.

The LabVIEW sub-VI element, included in the company supplied function library, can be used to read position and control movement throughout the indentation cycle. A LabVIEW indicator element can be used in conjunction with the provided sub-VI to obtain the current position of the piezo-stage. Additionally, this sub-VI can read a control element array that sends the stage a one-time command of how many micrometers to move. These reading and writing commands are encapsulated in a while-loop structure to discretely move the stage during the loading and unloading phases. To avoid lag, 10 Hz was chosen as the selected frequency for the commands.

When the LabVIEW program starts, position of the piezo-stage is read and reset it to its origin. Once the position is reset, the program shows a voltage-plot in real-time. This plot is used to adjust capacitive sensor using mini-stage (Fig. 1A-3) to ensure initial voltage is close to the upper limit of the capacitive sensor range so that indentations do not saturate the sensor. Additionally, the piezo-stage is reset in one displacement command (less than 10 ms), so the real-time voltage plot allows observation of environmental noise and ensures large-displacement-induced harmonic motion has dampened before beginning indentation.

#### II.A.1. Piezo-electric stage control for spring-loading

Real-time plots for piezo-stage displacement and indentation force are displayed for continual monitoring. During the loading phase, the code reads the desired indentation depth and indentation rate to move the piezo-stage. A time element is initiated outside the main while-loop to keep track of how much time has elapsed between displacement commands and acquired signals. Elapsed-time information (Δt_i_, s) is multiplied by the user-supplied indentation rate (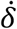, μm/s) to calculate the value of every displacement command (Δ*δ*, μm) that is sent to the piezostage during the loading phase:

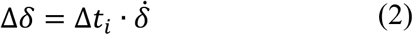

During the relaxation phase, the piezo-stage receives commands of zero displacement until total elapsed time surpasses the sum of loading and relaxation times. At that point, displacement commands are sent so that the piezo-stage returns to its initial position:

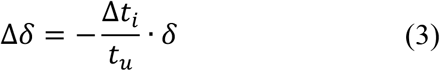

using unloading time (*t_u_*) input information.

At the end of the indentation, a force-displacement plot is displayed for the loading phase of the indentation with an exponential fit in the form of *F* = *C* · *δ^n^*. This exponent *n* of the loading curve is used to assess if indentation parameters, such as initial position and indentation depth, were properly selected for our chosen model. Clearly, this fitting routine and its parameters could be adjusted for other contact models of interest.

#### II.A.2. Piezo-electric stage control for stress-relaxation

True stress-relaxation differs from spring-loading in that it sends non-zero displacement commands to the piezo-stage to account for probe movement during the relaxation phase. To maintain constant indentation depth, displacement commands are sent to the piezo-stage to account for the additional sample indentation depth:

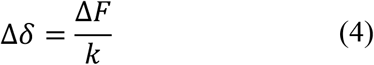

where Δ*F* is defined as the change in force between two consecutive displacement commands (i.e. the change in force for every Δ*t_i_*). Notice that Δ*δ* is negative, indicating movement away from the sample, because the change in force (Δ*F*) is negative as well.

During the unloading phase, displacement commands are sent so that the piezo-stage returns to its original position in the allotted time (unloading time, t_u_):

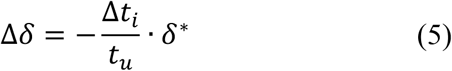

where *δ*\* is the piezo-stage position at the end of the relaxation phase, acquired by using the piezo-stage sub-VI indicator.

### II.B. Cantilever Design and Calibration

Our bi-layer, three-beam cantilevers are designed to minimize horizontal deflection with vertical deflections (Figure 3). Grade 5 titanium sheet was precision-cut into flexures via water jet (Micro Waterjet LLC, Figure 3A). The flexures are assembled into a single cantilever construct using a custom machined aluminum holder (Figure 3B-1), aluminum screws, and silver paint (Ted Pella, Inc., Leitsilber 200) to improve conductivity and maintenance of ground (Figure 3B-2). We used a carbon fiber rectangular shaft to connect the free ends of the cantilever without adding substantial weight and to enable modular indentation tip attachment (Figure 3B-3).

**Figure 3.**
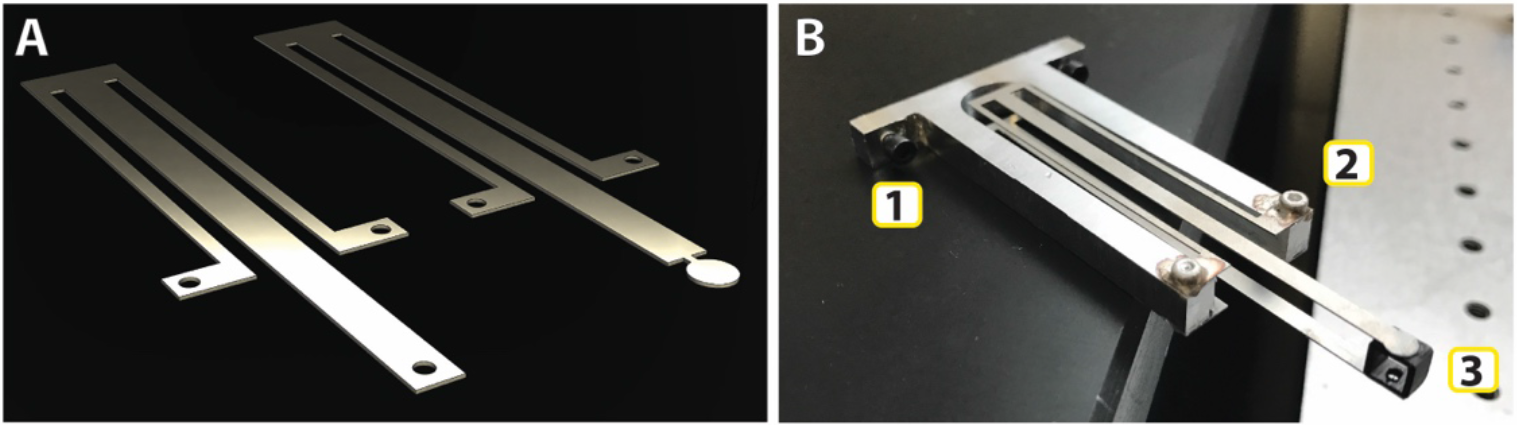
(A) General cantilever design for top (right) and bottom (left) flexures. (B) Assembled cantilever consists of (1) an aluminum holder that fastens to the aluminum plate (Figure 1A), (2) screws covered in metallic paint that electrically connect flexures, and (3) a carbon fiber rectangular shaft, cut and glued between flexures to minimize horizontal displacement.

Cantilever deflections are converted into forces using voltage of capacitive sensor and stiffness of cantilever. Cantilever deflection (*y_i_*) is calculated using measured Δ*V* and the relationship between distance and voltage from the capacitive sensors (e.g. sensor C8S-3.2-2.0, Section II):

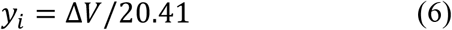

Then, Hooke’s Law is used to obtain indentation force (*F_i_*):

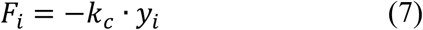

where *F_i_* is indentation force, *y_i_* is deflection from Equation 6, *k_c_* is cantilever stiffness.

Analytical calculations for cantilever stiffness yield values of *k_c_^soft^* ~ 20 N/m and *k_c_^stiff^* ~ 100 N/m using two different designs (example dimensions of soft cantilever in Supplementary Figure B), nominal values for Grade 5 titanium sheets (*E* = 110.3 GPa, t = 406 μm), and standard stiffness equations for cantilever beams (see example textbook^11^). Assembled cantilever stiffness is likely to vary from theoretical stiffness, so each assembled cantilever was calibrated using the capacitive sensor and small weights. Specifically, we weighed a segment of wire using a 0.01 mg resolution scale (Mettler Toledo XSE205 DualRange) and used the wire segment to load the free end of the cantilever, centered on the carbon fiber connector. Deflection was calculated by converting the measured change in voltage from the capacitive sensor to distance (Equation 6). The wire was then cut to make it lighter, weighed using the balance, and re-hung on the connector. Force, voltage, and deflection were recorded for 25 masses for each cantilever, which yielded *k_c_^soft^* = 19.63 ± 0.47 N/m (mean ± SD) using masses between 0.068 and 0.569 g and *k_c_^stiff^* = 79.8 ± 0.404 N/m using masses between 0.096 and 1.954 g.

### II.C. Indentation Probes

#### II.C.1. Custom diameter indentation tips

Borosilicate 1 mm diameter rods were used to fabricate hemispherical indentation tips with diameters ≤ 1 mm. Rods were heated and pulled using a glass micro-pipette puller (Narishige PP-830 Pipette Puller), and the resulting conical geometry was later heated by open flame to generate a smooth hemisphere. Temperature and load settings used to pull rod and reheat time in open flame affect final sphere diameter. Upright microscope was used to measure resulting spherical tips. These glass probes are then glued to an M2 screw (McMASTER-CARR, socket head, stainless steel) that screws into the 2 mm diameter hole in the bottom flexure of the cantilever (Figure 3A).

#### II.C.2. Polished ruby spheres

Three millimeter ruby ball lens and 4 mm ruby half-ball lens (Edmund Optics Worldwide, ruby-doped sapphire Al2O3) are glued to M2 screws and used as larger alternatives to our custom-made glass probes (mounted example in Figure 1A-6). Diameter tolarance is ±2.54 μm which is negligible in propagation of error calculations (see Section IIG). Also, the low surface roughness of optically polished ruby is supports assumptions of the popular Hertz model.

### II.D. Initiation of Indentation

Since accurate surface finding is critical to fit force-displacement data to contact models, a script that allows automated movement of the piezo-stage was included to position the probe in the correct location. The coarse manual stage is only used to visually bring the probe close the sample, after which the script can be used to finely tune the position of the piezo-stage and bring the probe into contact while reading the real-time voltage of the capacitive sensor. Once the probe is in position, the user interface allows the input of indentation depth, indentation rate, relaxation time, and unloading time (see Section II.A.). The front panel displays real-time displacement- and force-time plots that can be modified to show a smaller or larger time range in the x-axis.

Common soft matter test samples include hydrogels and *ex-vivo* tissue resections, which are typically submerged in culture medium or phosphate buffer saline (PBS) to keep them hydrated and reduce adhesion effects. To account for adhesion, common models such as Johnson-Kendal-Roberts (JKR)^12^ and Derjaguin-Muller-Toporov (DMT)^13^ can be used, though there are countless ways to describe mechanical properties of soft matter using indentation. Advanced methods and models are beyond the scope of this work, and the reader is directed to an exemplary review by Oyen and Cook.^14^

### II.E. Post-Processing

After the complete indentation cycle, a regression in the form of *F* = *C* · *δ^n^* is fit to force-displacement data from the loading phase. This fit allows the user to determine if the indentation was reasonable for their configuration, e.g., this coefficient should be close to 3/2 if using a spherical probe or 1 if using a flat punch (following Hertz model). Other coefficients might serve as an indication of a well-selected initial position for the indentation, e.g. n = 2 for indentations where surface area is greater than 10% the thickness of the sample.^10^ The final part of the code fits the force-displacement data during relaxation to the commonly used Standard Linear Solid model. However, the indentation data can be fit to any contact model and constitutive model the user considers relevant and applicable.

LabVIEW then generates *.csv files with time, piezo-stage position, sample indentation depth, and force. These files can be imported directly into the workspace of MATLAB to fit the loading and the relaxation portions of the indentation to different models. Additional functions enable batch processing and analysis, which is valuable for the large samples sizes needed for soft matter with high variability across samples.

### II.F. Treatment of Vibrational Noise

Though we attempt to isolate indentation equipment from environmental vibrations (TMC Isolation Table 63-543 HP), the cantilevered probe incurs vibrational noise from the piezo-stage during normal operation. During indentation, both the sensor and piezo-stage operate at 10 Hz, the maximum rate for the piezo-stage. Because the piezo-stage is receiving only 10 instructions per second, the displacement per instruction can be large for a given displacement rate which causes the cantilever tip to vibrate. This vibration is captured by the capacitive sensor and appears to amplify noise of recording.

This vibration can be both a strength and a weakness. Additional noise in the voltage signal, which is later used to calculate forces, can hinder data fitting. However, since high vibration is only present when the tip of the cantilever is free to move, i.e. away from contact with a solid surface, a *reduction* in vibrational noise can indicate contact with sample surface (Figure 4). This reduction in noise upon first contact can be particularly helpful for soft matter samples that are typically submerged to reduce adhesion effects. Probe displacement through fluid generates quantifiable buoyancy forces that could be confounded with indentation forces during surface finding, making reduction in noise due to cantilever vibration helpful to identify position of first contact.

**Figure 4.**
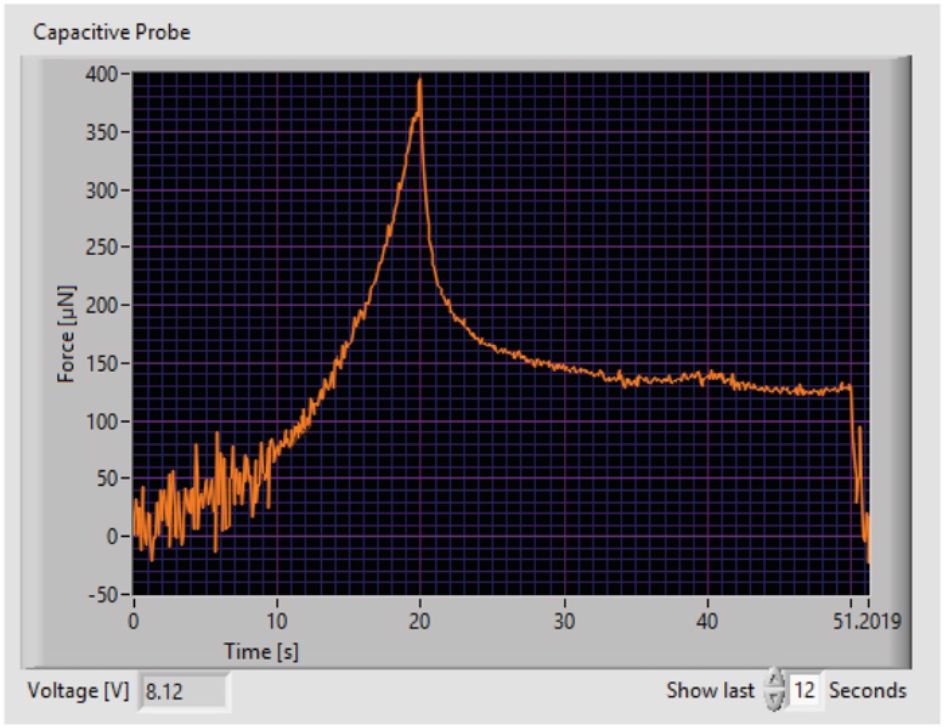
Force vs. time plot shows high noise before contact with the sample (~10 s), presenting a predictable indicator for initial contact. Because the stage is displaced at a user defined indentation rate, distance between the original position of the stage and the sample can be calculated. This information can be used to move the piezo-stage before beginning the next indentation cycle.

### II.G. Sources and Propagation of Uncertainty

As a case study, we considered the uncertainty propagation for an example contact model. End users are not constrained to this model, though; see discussion of contact and viscoelastic models in example review paper.^15^ Here, we use a rearrangement of the common Hertz contact model between a sphere (the probe) and half space (the hydrogel) to determine the elastic modulus at a given point in time:^3,16,17^

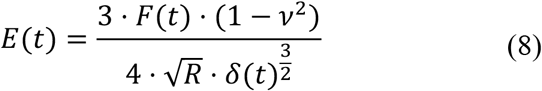

where E(t) = total elastic modulus as a function of time, F(t) = force calculated from deflection of cantilever as detected by the capacitive sensor; *ν* = Poisson’s ratio; R = radius of indenter tip; and *δ*(*t*) = indentation depth of cantilever tip.

The resolution of the capacitive sensor and its associated error propagation are small compared to the force fluctuation caused by the vibration of the cantilever during the loading or unloading phases. Propagation of uncertainty (see example textbook^18^) is carried out using estimated error values for force, Poisson’s ratio, and displacement (Table 1). While propagation analysis shows that percentage error values can reach around 25% for indentation parameters that yield low forces throughout indentation (Figure 5), 25% overall error is still reasonable compared to the large differences anticipated between groups (e.g. healthy vs. diseased tissue).

**Table 1.**
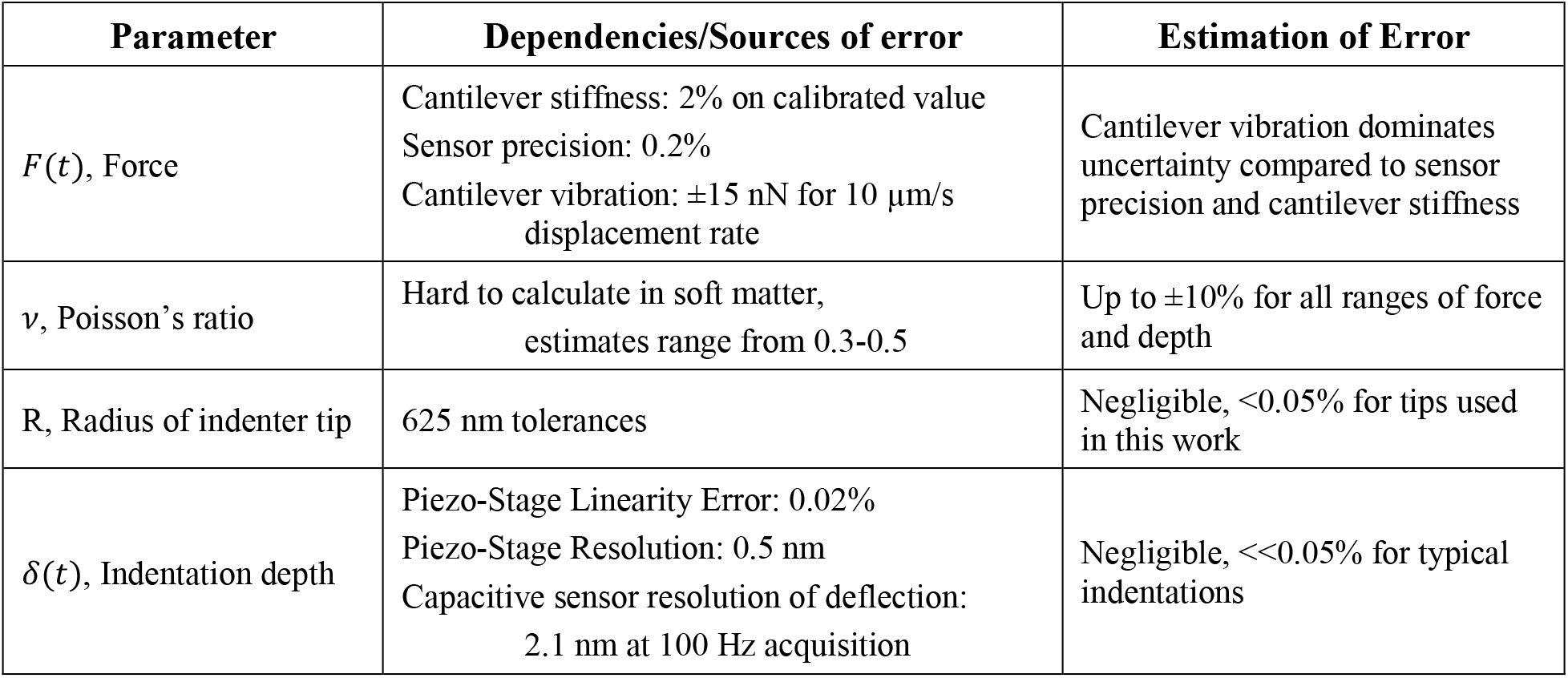
Uncertainty on metrics used to calculate example material property, transient modulus, from Equation 8 as visualized in Figure 5.

**Figure 5.**
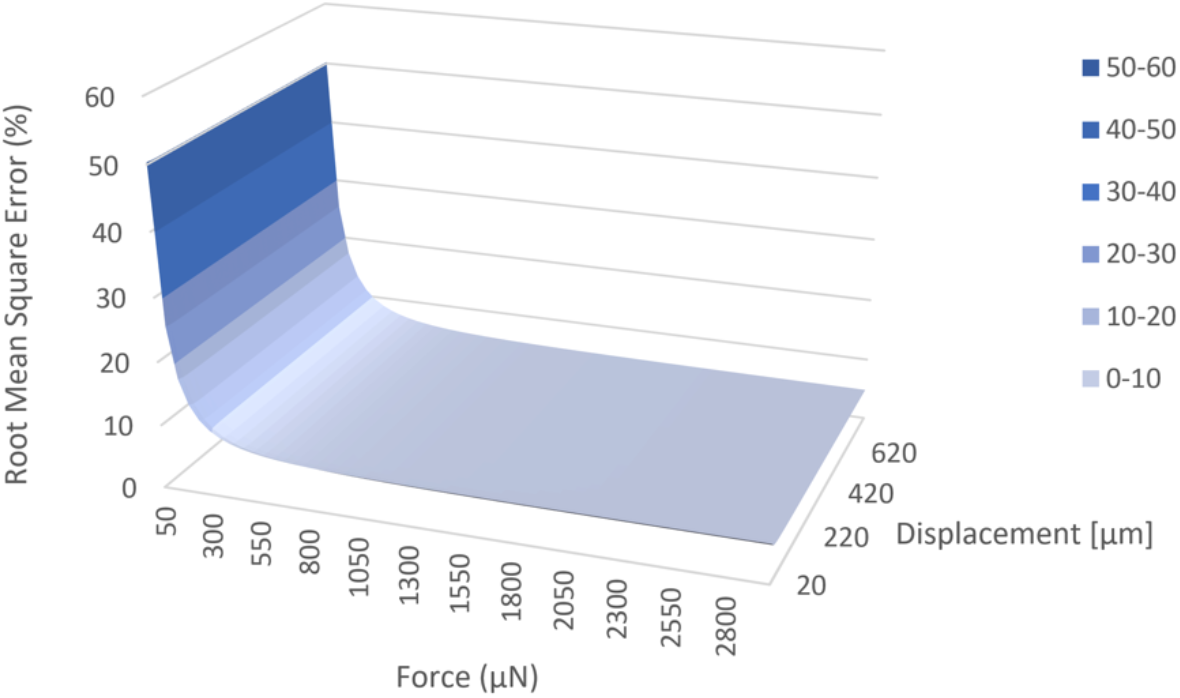
Error propagation calculations for values of displacement and force within the system’s ranges show high dependency on force values. For reference, error propagation for soft collagen hydrogels (850 Pa: 100 μN and 140 μm) yields 25% error. Indentation rate of 10 μm/s was used for all indentations.

## III. EXPERIMENTAL DEMONSTRATION

A suite of hydrogels and silicones were indented with the system to show measured stiffness values are found within those reported in the literature (Figure 6A). Samples were stored in water and indented no less than 3 hours after fabrication (typically 24-48 hours after fabrication) to allow swelling to reach a steady-state and not interfere with acquired data during indentation.

**Figure 6.**
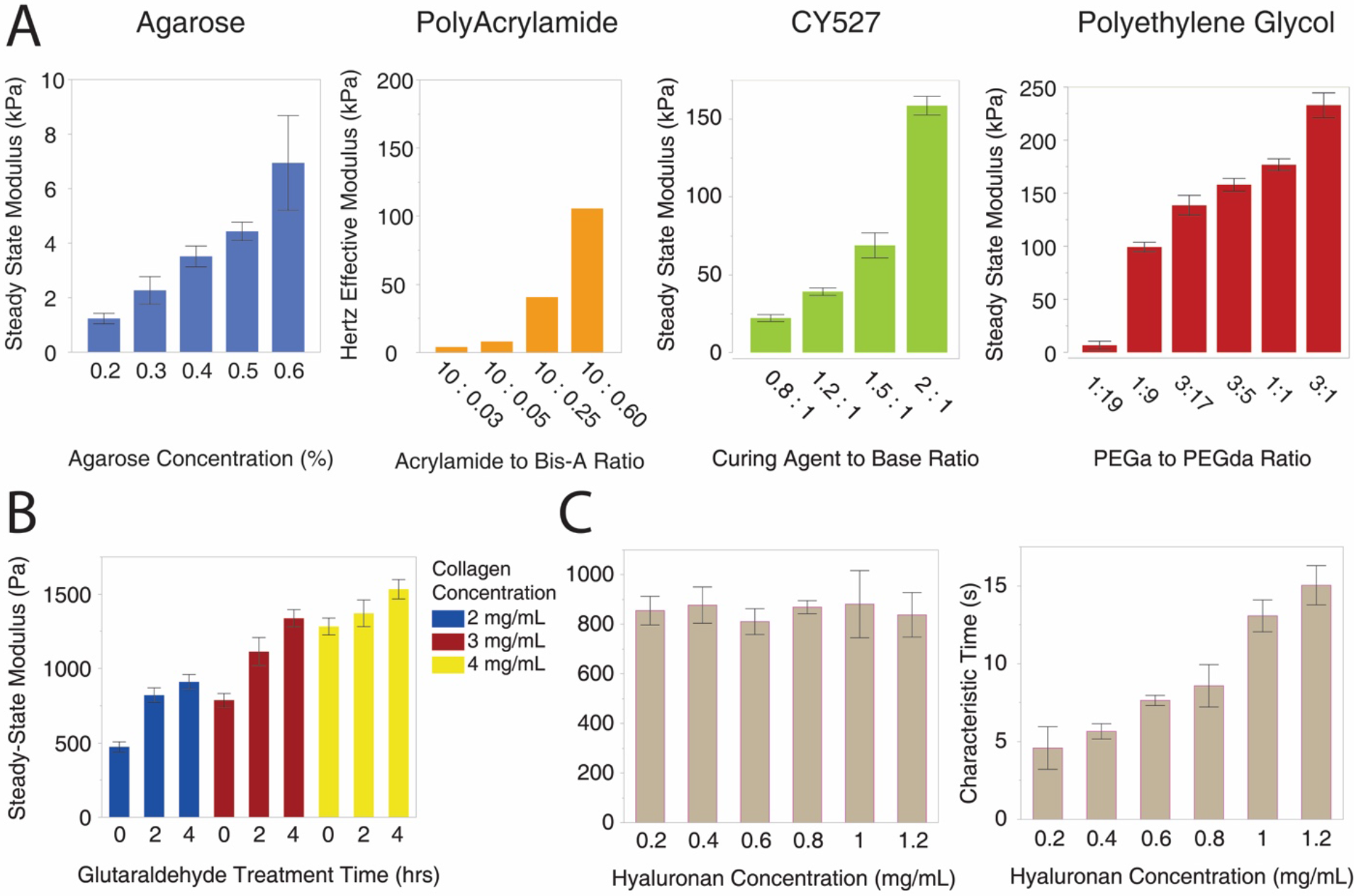
(A) Soft hydrogels, quantified by our system described here, have SSM ranging two orders of magnitude. See Section III for experimental details. (B) SSM results for low concentration collagen hydrogels at different concentrations for different cross-linking periods, showing the capabilities of our system to quantify subtly changing mechanical properties in the sub-kPa range. (C) Characteristic SSM and relaxation times for 3 mg/mL collagen gels show that our system can also identify changes in viscous properties while SSM stays constant due to varied hyaluronan concentrations.

Agarose powder (UltraPureTM Agarose, Life Technologies Corporation) was diluted in water, heated to approximately 70°C, and stirred for 15 minutes until the solution became transparent. Solution is transferred to petri dishes and allowed to cool to achieve ~3 mm-thick agarose hydrogels. After 30 minutes of cooling at room temperature, water is added to the petri dish to fully submerge the sample and maintain hydration.

Polyacrylamide (PA) gels were polymerized from acrylamide-bisacrylamide precursor solution with ammonium persulfate and tetramethylethylenediamine initiators, and gel volumes of 380 μL were covered with 22 mm-diameter cover glass during polymerization, which resulted in ~1 mm-thick discs of gel.

Silicone CY527 (Dow Corning) was fabricated following manufacturer’s instructions. CY527 is a platinum-based silicone cure system fabricated by combining Part A (platinum complex) and part B (vinyl-functional siloxane) in A:B ratios of 0.8:1, 1.2:1, 1.5:1, and 2:1 to produce silicone samples of varied stiffness. Precursor solution was thoroughly mixed, poured into the bottom of a 40-mm diameter petri dish, vacuum degassed for an hour, and kept in a 50°C oven overnight. Detergent-spiked water was used to reduce adhesiveness and keep the samples submerged during indentation.

Polyethylene glycol (PEG) methyl ether acrylate and PEG diacrylate were combined in 95:5 and 90:10 weight to weight ratios. Total polymer of 25 wt% in water was vortexed, and Irgacure (2-Hydroxy-4’-(2-hydroxyethoxy)-2-methilpropiophenone) was added and used as the photo-initiator. Volumes were poured into petri dishes and subjected to UV to achieve 3 mm thick polymerized samples.

Collagen samples were fabricated using a variety of methods to change expected mechanical properties. For the basic protocol, high concentration rat tail collagen type I (Corning) was diluted with 0.02% acetic acid and combined in a 3:1 ratio with Dulbecco’s Modified Eagle’s Medium 5x (SIGMA Life Science) and 1M HEPES buffer solution (Gibco by Life Technologies) to fabricate 2, 3, and 4 mg/mL collagen hydrogels. Precursor solution was prepared at ~4°C and then incubated at 37°C for 30 minutes to allow thermogelling. Gels were then hydrated with PBS and kept at 37°C until before indentation (between 2-24 hours). Replicate gels were treated gluteraldehyde to increase cross-linking and therefore stiffness (Figure 6B). For hyaluronic acid (HA)-containing gels, HA sodium salt from *Streptococcus equi* (Sigma-Aldrich) was dissolved in distilled, sterile H2O (5mg/mL dilution) and added to collagen gel precursor solution. Collagen-HA composite was then allowed to thermogel at 37°C for 35 minutes before adding cell culture medium (Figure 6C).

Using Equations 1 and 8, the total elastic modulus as a function of time, E(t), was calculated for tip radius R = 2.0 mm. A typical assumption for soft matter and biological tissue is that it is incompressible, i.e. Poisson’s ratio, *ν*, = 0.5. Next, an appropriate constitutive model must be fit to the adjusted force-displacement data. The most basic constitutive models of engineering materials are linear elastic, but these typically are inadequate to describe liquid-containing soft matter like hydrogels and tissue. Viscoelastic models are often more appropriate for biological tissues, combining linear elastic components with elements that represent polymer strands relaxing and interacting. The standard linear solid (SLS) is a common viscoelastic mechanical network model, composed of two springs representing linear elasticity and one dashpot representing viscosity. To determine the parameters of the SLS model for a given sample, we fit the relaxation portion of the total elastic modulus curve, *E*(*t*), to the following exponential function:

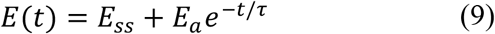

where *E_ss_* = the Steady-State Modulus (SSM) after relaxation, *E_a_* = the additional or strain-rate dependent modulus, *η* = viscosity, and τ = *E_a_/η*

The strain-rate-dependent and “liquid-like” properties of the material are represented by viscosity, η, and as the material relaxes, the exponential multiplying *E_a_* trends to zero. The resulting *E_ss_* can be interpreted as “steady-state modulus” (SSM), also known as quasi-static modulus, infinity modulus, aggregate modulus, and/or relaxation modulus. This simple combination of linear elastic and viscous components has an easily understood physical interpretation, making this model particularly useful for interdisciplinary investigations with collaborators from broad backgrounds.

Hertz and SLS fits are achieved using Non-Linear Least Squares fitting and the Levenberg-Marquardt algorithm as a Custom Equation fit-type in MATLAB’s regression toolbox. While we chose Hertz and SLS to interpret force-displacement data, we reiterate that this indentation system is compatible with any contact and constitutive models researchers deem appropriate for their samples.

With this suite of soft matter characterization, we demonstrate that our system is capable of calculating elastic and time-dependent properties of soft matter for a wide range of stiffness values. The ability of our system to work with different sensors, probes, and cantilevers makes it an ideal characterization tool for soft matter used in a wide range of applications, including biomedical and tissue engineering.

## IV. DISCUSSION AND FUTURE APPLICATIONS

The wide variety of studies attempting to characterize mechanical properties of biological tissue and soft matter underscore the challenge of soft matter characterization, and the high variability in reported metrics and techniques reinforce the complexity in choosing and applying appropriate models to soft matter. Soft materials (100 Pa – 1 MPa) with highly irregular geometries and sizes – *ex-vivo* resections, especially – violate some or many of the physical assumptions of existing contact models that work well for stiffer and more homogeneous materials and structures.

While development of complex constitutive models appropriate for computational applications will require extensive mechanical characterization, mesoscale indentation can help researchers in biomedical fields identify simple design parameters for tissue engineering applications. Our group, for example, has used this device to quantify mechanical properties of diseased and healthy tissues of animals and patients. We have shown that stem cell therapy can restore elasticity to myocardium tissue of hypertensive rats, making use of the millimetric scale of the system’s indentation probes to differentiate between left and right ventricle walls.^2^ Our group has also shown that there are correlations among inflammatory bowel disease, fibrosis, and stiffness in humans^17^ and hypertension-linked gut dysbiosis, gut fibrosis and stiffness in rats.^19^ We have also used this system to compare native tissue to reconstituted implantable hydrogels.^20^

As discussed, indentation is a convenient method for researchers with limited access to sophisticated equipment and limited experience in experimental mechanics, but models leveraging indentation to describe stresses and strains in heterogeneous, anisotropic, highly-adhesive, and soft materials remains challenging. Mesoscale indentation systems, like the one we describe here, can facilitate researchers interested in matching primary tissue mechanics to synthetic and engineered scaffolds by enabling simplifying assumptions as well as support researchers interested in developing advanced contact and constitutive models.

## V. ACKNOWLEDGMENTS

The authors gratefully acknowledge support from NIH R01HL102033, NIH R01HL132448, NSF BMMB 1636007, and Medtronic AGR DTD 07-28-2015. We also thank Benjamin Stadnick, Richard Nay, Syed Asif, Rajiv Dama, and Thomas Wyrobek of Hysitron-Bruker for helpful technical discussions.

